# Water availability drives signatures of local adaptation in whitebark pine (*Pinus albicaulis* Englm.) across fine spatial scales of the Lake Tahoe Basin, USA

**DOI:** 10.1101/056317

**Authors:** Brandon M. Lind, Christopher J Friedline, Jill L. Wegrzyn, Patricia E. Maloney, Detlev R. Vogler, David B. Neale, Andrew J. Eckert

**Author notes:** Corresponding Author. Brandon M. Lind, 1000 West Cary Street, Life Science Bldg. - Department of Biology Rm 126, Virginia Commonwealth University, Richmond, Virginia 23284 USA. Current Address: Department of Biology, Virginia Commonwealth University, Richmond, VA 23284.

## Abstract

Patterns of local adaptation at fine spatial scales are central to understanding how evolution proceeds, and are essential to the effective management of economically and ecologically important forest tree species. Here, we employ single and multilocus analyses of genetic data (*n* = 116,231 SNPs) to describe signatures of fine-scale adaptation within eight whitebark pine (*Pinus albicaulis* Engelm.) populations across the local extent of the environmentally heterogeneous Lake Tahoe Basin, USA. We show that despite highly shared genetic variation (*F*_ST_ = 0.0069) there is strong evidence for adaptation to the rain shadow experienced across the eastern Sierra Nevada. Specifically, we build upon evidence from a common garden study and find that allele frequencies of loci associated with four phenotypes (mean = 236 SNPs), 18 environmental variables (mean = 99 SNPs), and those detected through genetic differentiation (*n* = 110 SNPs) exhibit significantly higher signals of selection (covariance of allele frequencies) than could be expected to arise, given the data. We also provide evidence that this covariance tracks environmental measures related to soil water availability through subtle allele frequency shifts across populations. Our results replicate empirical support for theoretical expectations of local adaptation for populations exhibiting strong gene flow and high selective pressures, and suggest that ongoing adaptation of many *P. albicaulis* populations within the Lake Tahoe Basin will not be constrained by the lack of genetic variation. Even so, some populations exhibit low levels of heritability for the traits presumed to be related to fitness. These instances could be used to prioritize management to maintain adaptive potential. Overall, we suggest that established practices regarding whitebark pine conservation be maintained, with the additional context of fine-scale adaptation.

## Introduction

The study of local adaptation has been an integral part of evolutionary biology as a whole, as local adaptation influences a wide variety of biological patterns and processes (reviewed in Savolainen *et al*. 2013). Trees in particular have received much attention in this regard because many species are ecologically and economically important, and high outcrossing rates (Neale & Savolainen 2004) result in large effective population sizes (which increase the effectiveness of selection) as well as weak neutral genetic differentiation (which decreases the confounding effects of selection and population structure), which together create ideal circumstances in which to detect selective processes in nature (Savolainen & Pyhäjärvi 2007). Investigators seeking to explain the genetic basis of local adaptation in trees, and plants in general, have been motivated by observations of significant differentiation for quantitative genetic variation across populations (e.g., *Q*_ST_). If such phenotypes have a genetic basis, the underlying loci may be differentiated among populations as well (Endler 1977; reviewed in Storz 2005, Haasl & Payseur 2016). In these cases, loci contributing to local adaptation could be identified through genetic indices of differentiation, or by targeting trait- or environmentally-associated loci that stand out above background demography. Yet, theoretical (Latta 2003; Le Corre & Kremer 2003) and empirical (Hall *et al*. 2007; Luquez *et al*. 2007) investigations have shown that discordance between *Q*_ST_ and *F*_ST_ of causative loci can occur under adaptive evolution. Moreover, as the number of underlying loci increases, the divergence between these indices increases as well, and the contribution of *F*_ST_ to any individual underlying locus decreases. In cases that exhibit strong diversifying selection and high gene flow, this adaptive divergence results from selection on segregating genetic variation (Hermisson & Pennings 2005; Barret & Schluter 2008) and is attributable to the among-population component of linkage disequilibrium (Ohta 1982, Latta 1998). In the short term, local adaptation will be realized through subtle coordinated shifts of allele frequencies across populations causing covariance (i.e., LD) among many underlying loci (Latta 1998; Barton 1999; Latta 2003; McKay & Latta 2002; Kremer & Le Corre 2012; Le Corre & Kremer 2012), such that adaptation need not take place through numerous fixation events or sweeping allele frequency changes (MacKay *et al*. 2009; Pritchard & di Rienzo 2010). Over many thousands of generations, these shifts can lead to concentrated architectures of large-effect loci with a reduction of those with small effect (Yeaman & Whitlock 2011). For studies investigating continuous phenotypes such as those often related to fitness, even among populations with highly differentiated phenotypic traits sampled under a robust design (Lotterhos & Whitlock 2015), it may be difficult to identify many of the loci underlying the quantitative trait in question. Thus, for many species, specifically across fine spatial scales, the signal of local adaptation within much of current genetic data may go largely undetected using only single-locus approaches (Latta 1998; 2003; Le Corre & Kremer 2003; Yeaman & Whitlock 2011; Kemper *et al*. 2014), resulting in calls for theory and empiricism that move beyond single-locus perspectives (Pritchard & di Rienzo 2010; Sork *et al*. 2013; Tiffin & Ross-Ibarra 2014; Stephan 2015).

Populations of forest trees, particularly conifers, have a rich history of common garden, provenance tests, and genecological studies that demonstrate abundant evidence for local adaptation among populations, even over short geographic distances (e.g., Mitton 1989; 1999; Budde *et al*. 2014; Csilléry et al. 2014; Vizcaíno *et al*. 2014; Eckert *et al*. 2015) providing further support that fine spatial scales are relevant to adaptation (Richardson *et al*. 2014). This extensive history has also revealed the highly polygenic nature of adaptive traits (Langlet 1971; Holland 2007). Even so, the majority of these investigations have been limited to single-locus perspectives using either candidate genes (e.g., González-Martínez *et al*. 2008; Eckert *et al*. 2009) or a large set of molecular markers (e.g., Eckert *et al*. 2010) to explain the genetic basis of local adaptation. In most cases, a few loci underlying the adaptive trait in question are identified and generally explain a small to moderate proportion of the overall heritability of the trait (Neale & Savolainen 2004; Savolainen *et al*. 2007; Ćalić *et al*. 2016). Yet because of the presumed polygenic nature underlying these adaptive phenotypic traits, and because past investigations have generally applied single-locus perspectives, it is likely that a majority of the genetic architecture of local adaptation in trees remains undescribed (Savolainen 2007; Sork *et al*. 2013; Ćalić *et al*. 2016).

Spurred in part by the advance of theory and availability of genome-wide marker data, attention has been refocused to describe underlying genetic architectures from a polygenic perspective. This transition began in model organisms (e.g., Turchin *et al*. 2012) and has expanded to other taxa such as stick insects (Comeault *et al*. 2014; 2015), salmon (Bourret *et al*. 2014), and trees (Ma *et al*. 2010; Csilléry *et al*. 2014; Hornoy *et al*. 2015). Indeed, species that occupy landscapes with high degrees of environmental heterogeneity offer exemplary cases with which to investigate local adaptation. Near its southern range limit, whitebark pine (*Pinus albicaulis* Engelm.) populations of the Lake Tahoe Basin (LTB) inhabit a diversity of environmental conditions. As exemplified by the strong west to east precipitation gradient (see Figure 1), many of the environmental characteristics of the LTB vary over short physical distances (<1km) and have the potential to shape geographic distributions of *P. albicaulis* at spatial scales below those typically investigated (i.e., range-wide studies) for forest trees. Local spatial scales are of particular interest to resource and conservation agencies as this is the scale at which most management is applied. Here, we build upon past work from a common garden (Maloney *et al*. in review) to investigate the genetic architecture of fine-scale local adaptation across *P. albicaulis* populations of the LTB by exploring the relationships between genotype, 18 environmental variables, and five fitness-related phenotypic traits using both single and multilocus approaches. Specifically, we use the *P. albicaulis* populations of the LTB to address the following four questions: (i) Is there evidence that a long-lived, outcrossing plant species exhibiting high levels of gene flow can be locally adapted across fine spatial scales? (ii) What is the genetic basis and relationship among loci underlying adaptation in such a species? (iii) How similar are the genetic bases of fitness-related phenotypes to the loci putatively under selection from the environment? Using this information, we will contextualize how instances of fine-scale adaptation have management implications. This study highlights the advantages of a polygenic perspective and investigates signatures of local adaptation using a large set of null markers to determine the extremity of allele covariance among putatively adaptive loci where others have relied on simulation or null candidate genes. Furthermore, this work provides additional empirical evidence for theoretical predictions of covariance among adaptive loci found by other studies in trees.

**Figure 1.**
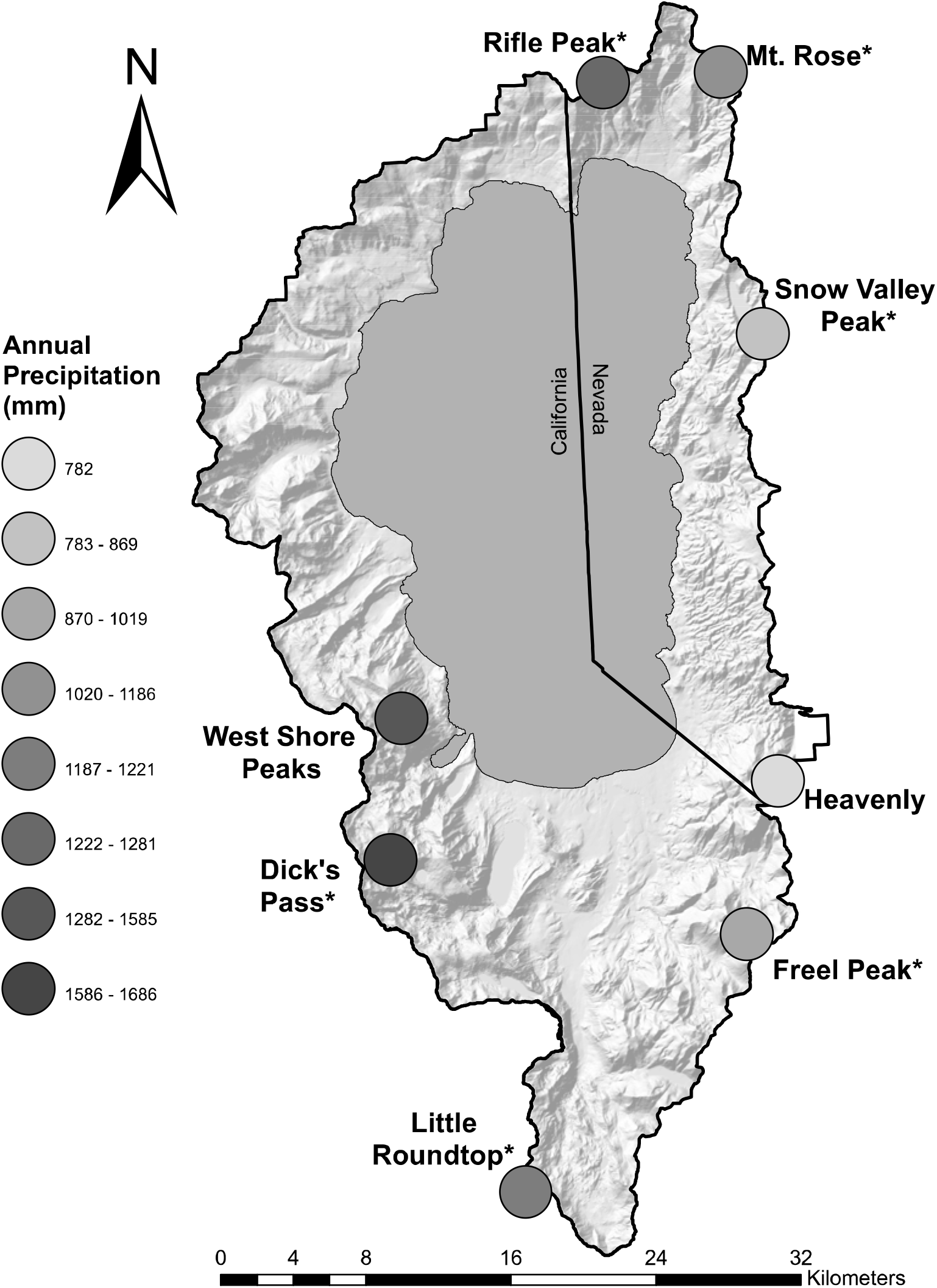
Populations used for sampling P. albicaulis within the Lake Tahoe Basin (dark outline). Annual precipitation is given for each population to demonstrate the west-east rain shadow experienced across fine spatial scales. Asterisks indicate populations in the common garden study.

## Materials and Methods

### Focal species, study area, and sampling

A principal component of high elevation forests in California and Nevada, *P. albicaulis* is widespread throughout subalpine and treeline environments and plays a vital role in ecosystem function and services including food resources for wildlife, forest cover, watershed protection, protracting snowmelt, and biodiversity (Hutchins & Lanner 1982; Farnes 1990; Tomback *et al*., 2001; McKinney *et al*., 2009; Tomback & Achuff 2010; Tomback *et al*. 2016). Most of the species' distribution is outside of California, extending northward into Oregon, Washington, British Columbia, and Alberta and eastward into northern Nevada, Idaho, Montana, and Wyoming (Critchfield & Little 1966; Tomback & Achuff 2010). Whitebark pine is a foundation species in subalpine ecosystems throughout most of its range in western North America (Ellison *et al*. 2005) and is threatened by fire-suppression, climate change, the non-native pathogen white pine blister rust, caused by *Cronartium ribicola* J.C. Fisch., and mountain pine beetle, *Dendroctonous ponderosae* Hopkins (Tomback & Achuff 2010; Mahalovich & Stritch 2013).

The LTB lies within California and Nevada in the north-central Sierra Nevada range, varies in elevation from 1900 to 3300m, and is flanked to the west by the Sierra Nevada crest and to the east by the Carson Range. The LTB experiences a Mediterranean climate with warm, dry summers and cool, wet winters. Precipitation falls during the winter months, most often in the form of snow, with a strong west-east gradient. The geology of the region is dominated by igneous intrusive rocks, typically granodiorite, and igneous extrusive rocks, typically andesitic lahar, with small amounts of metamorphic rock (USDA NRCS 2007).

Each of the eight study populations (three subplots per population) were located in a distinct watershed and distributed around the Basin to capture variation in the physical environment (e.g., climate, geology, and topography; Figure 1). Needle tissue was sampled in the summer of 2008 from 244 *P. albicaulis* trees (Table 1). From these eight populations, six populations were chosen to sample cones from 88 of the trees that were sampled for needle tissue. All samples were collected from trees separated by 30 to 1000m, with an average interpopulation distance of 31 km. Universal Transverse Mercator coordinates, elevation, slope, and aspect (USDA FS FHTET) were used with the PRISM climatic model (Daly *et al*. 1994) to determine climatic parameters of sampled areas from 1971-2000, while soil survey data (USDA NRCS 2007) were used to describe the edaphic conditions of the LTB (Table 1).

**Table 1.** *Population location and associated attributes. Population size* – total (maternal trees with seedlings in common garden). *Climatic values were ascertained from data spanning 1971-2000. Ann. precipitation – annual precipitation; AWC – available water capacity at 25cm or 50cm soil depth; CEC – cation exchange capacity; GDD – growing degree days above 5°C; Max solar rad input – maximum solar radiation input; WC-15bar-water capacity at-15bar (wilting point); WC-⅓bar – water capacity at-⅓bar (field capacity). Asterisks indicates populations from which seeds sampled from cones were planted in a common garden. Environmental variables are averaged across subplots.*

|  | Dick's Pass* | Freel Peak* | Heavenly | Little Round Top* | Mt. Rose Ophir* | Rifle Peak* | Snow Valley Peak* | West Shore Peaks |
| --- | --- | --- | --- | --- | --- | --- | --- | --- |
| Population size | 25 (15) | 48 (19) | 25 (0) | 25 (14) | 49 (11) | 24 (15) | 24 (14) | 24 (0) |
| Ann. precipitation (mm) | 1686 | 1019 | 782 | 1221 | 1186 | 1281 | 869 | 1585 |
| AWC-25cm (kPa) | 1.66 | 1.57 | 1.12 | 1.97 | 1.95 | 1.89 | 2.66 | 1.20 |
| AWC-50cm (kPa) | 2.75 | 2.38 | 2.00 | 2.93 | 2.75 | 3.11 | 4.22 | 2.02 |
| CEC (cmol<sub>c</sub> · kg<sup>-1</sup>) | 0.00 | 1.45 | 0.00 | 12.50 | 2.90 | 0.00 | 0.00 | 0.00 |
| Clay (%) | 6.50 | 4.50 | 6.70 | 14.60 | 3.00 | 6.75 | 6.80 | 6.00 |
| Elevation (m) | 2806 | 2865 | 2851 | 2875 | 2717 | 2819 | 2740 | 2780 |
| GDD Aug (days) | 295 | 190 | 276 | 211 | 296 | 235 | 289 | 279.5 |
| GDD May (days) | 0 | 0 | 6 | 0 | 11 | 0 | 2 | 1 |
| Max solar rad input (%) | 83.59 | 79.03 | 78.40 | 80.09 | 90.61 | 93.28 | 71.70 | 76.43 |
| Max Temp – July (°C) | 21.1 | 21.6 | 23.2 | 21.5 | 22.9 | 22.7 | 23.4 | 21.8 |
| Min. Temp – Jan (°C) | -6.5 | -8.8 | -7.5 | -8.0 | -7.4 | -7.4 | -7.7 | -6.6 |
| Rock coverage (%) | 31.00 | 18.67 | 25.00 | 14.67 | 7.00 | 30.00 | 26.67 | 42.67 |
| Sand (%) | 77.67 | 87.80 | 83.50 | 66.20 | 90.60 | 74.00 | 64.50 | 85.00 |
| Silt (%) | 15.8 | 7.7 | 9.7 | 19.1 | 6.4 | 19.2 | 28.7 | 9.0 |
| WC-15 bar (kPa) | 6.6 | 4.0 | 3.3 | 3.6 | 5.5 | 8.7 | 14.0 | 2.5 |
| WC-1/3 bar (kPa) | 9.7 | 8.0 | 8.4 | 7.3 | 9.8 | 11.4 | 14.4 | 6.2 |

### Common gardens and phenotypic measurements

Fitness-related traits related to survival, especially during seedling and juvenile stages, are an important component of total lifetime fitness (e.g., Postma & Âgren 2016), particularly for forest trees, and are likely to be composed of phenotypic traits related to growth, phenology, resource allocation patterns, water-use efficiency, and disease susceptibility. In order to estimate early-lifetime phenotypes of mother trees, seeds sampled from 11 to 19 maternal trees (*n* = 88) located in six of the eight populations were established in a common garden (Table 1) using a random block design (for further details see Maloney *et al*. in review). Growth (height), phenology (budset), water-use efficiency (δ^13^C), and resource allocation [root:shoot biomass, N(μg)] were measured when seedlings reached ~2 years in age (see Maloney *et al*. in review for details). Height was recorded in April and October 2011, while 2 seedlings per family per block were harvested, clipped above the root collar, dried, and weighed to determine root and shoot biomass. For δ^13^C and N(μg) analysis, needle tissue from 1 seedling per family per block was harvested, coarsely ground, and dried at 60°C for 96 hours. Between 2-3mg of tissue per sample was sent to the Stable Isotope Facility at UC Davis for isotope analyses (http://stableisotopefacility.ucdavis.edu/).

### DNA extraction, sequencing, and analysis

Total genomic DNA was isolated from finely ground needle tissue sampled from 244 trees across all eight populations using the Qiagen DNEasy 96 Plant kit according to protocol (Qiagen, Germantown, MD). Restriction site-associated double digests of total genomic DNA using MseI and EcoRI enzymes (ddRADSeq, Peterson *et al*. 2012) were used to prepare three multiplexed libraries of up to 96 individuals each, as in Parchman *et al*. (2012). Total genomic DNA was digested and ligated with adapters containing amplification and sequencing primers as well as barcodes for multiplexing. These 96 barcodes (Parchman *et al*. 2012) are of 8-10bp sequences that differ by at least four bases. Ligated DNA was then amplified using high-fidelity PCR. Using the QIAquick Gel Extraction Kit (Qiagen), amplified fragments were then isolated near 400bp of pooled PCR product separated in a 1% agarose gel at 100V for one hour. Single-end sequencing of libraries was carried out on the Illumina HiSeq 2500 platform with a single library per flowcell lane. For added coverage, each library was sequenced twice using 50bp reads and twice for 150bp reads, except Library 3 which was sequenced 4x for 150bp reads to increase optimality of the mapping reference individual. All sequencing was performed at the DNA Sequencing Facility of the University of California at Berkeley (https://mcb.berkeley.edu/barker/dnaseq/home). After calling genotypes, SNPs, and further filtering (see Supporting Information), we judged the veracity of our sequence data by mapping the empirical set of SNPs against the sugar pine (*P. lambertiana* Dougl.) reference genome (v1.0) using 85% similarity and 50 length coverage thresholds (http://dendrome.ucdavis.edu/ftp/GenomeData/genome/pinerefseq/Pila/v1.0/).

### Identifying focal sets of loci

To identify genotype-environmental associations, we implemented bayenv2 (v2.0; Coop *et al*. 2010; Günther & Coop 2013), a Bayesian single-locus approach that accounts for population history and gene flow before performing association analysis (Coop *et al*. 2010). To ensure convergence, we ran five independent chains of bayenv2 using the empirical SNPs (n = 116,231), with 100,000 iterations for each SNP within each chain. MCMC convergence across chains was inspected using the coda library in R. For each SNP, we calculated the harmonic mean across chains for the Bayes factor (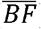) and absolute value of Spearman's ρ (hereafter 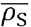). When calculating 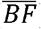, if a particular SNP returned Bayes factors greater than one for at least 3/5 chains, we would take the harmonic mean from this subset to avoid underestimation of the Bayes factor. However, if this was not the case (BF > 1 in ≤ 2/5 chains) we took the harmonic mean from the values that were less than or equal to one. We identified focal SNPs by the intersection between the upper tail (99.5^th^ percentile) of 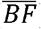 and the upper tail (99^th^ percentile) of the absolute value of 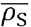, as recommended in the bayenv2 manual (v2.0; page 4).

To associate genotype with phenotype, we implemented a Bayesian sparse linear mixed model (BSLMM) from the GEMMA software package (Zhou *et al*. 2013). BSLMM is a hybrid of LMM and Bayesian variable selection regression (BVSR) that also offers considerable statistical advantages over single-locus GWAS approaches (Guan & Stephens 2011; Ehret *et al*. 2012; Zhou *et al*. 2013; Moser *et al*. 2015). Specifically, to describe the underlying genetic architecture, BSLMM uses priors (described below) and attributes of the genetic data to estimate the number of underlying SNPs (N_SNP_), the posterior inclusion probability (γ, hereafter *PIP*) for individual SNPs as well as the proportion of phenotypic variance explained by the polygenic and sparse effects of each SNP (*PVE*).

Before input to GEMMA, the empirical set of SNPs was reduced to include only those individuals with seedlings in the common garden (n = 88), and loci which had MAF below 0.01 due to this reduction were eliminated alongside monomorphic SNPs. For each phenotype, we ran four independent chains for the BSLMM, with 1,000,000 warm-up steps and 50,000,000 steps in the MCMC, sampled every 1000^th^ step. Priors for *PVE* by the model, *h*, were set as [0.01,0.9], and the log_10_ inverse number of SNPs, log_10_(l/p), [-3.0,0.0], which equates to between 1 and 300 underlying loci (*N*_SNP_). Convergence of the MCMC across chains was inspected using the coda library in R. To summarize the GEMMA output, we report means and 95 credible intervals for and from the posterior distributions. To assess significance of association of a SNP to a phenotype, we used the *PIP* from all four independent chains to calculate the harmonic mean (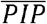) and chose SNPs that were greater than or equal to the 99.9^th^ percentile of 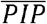 (n ~ 116 SNPs/phenotype) for each phenotype. We also explored SNPs with 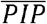 ≥99.8^th^ percentile (n ~ 232/phenotype).

We implemented the program OutFLANK (Whitlock & Lotterhos 2015) to investigate the opportunity of detecting the environmentally- and phenotypically-associated loci using outlier approaches based on population genetic structure alone (e.g., *F*_ST_). Using this approach and excluding loci with expected heterozygosity values below 10% with subsequent trimming of the lower and upper 5% of empirical *F*_ST_ values, we inferred a null distribution of *F*′_T_ and identified outlier loci with a false discovery rate of 5 from the empirical set of SNPs.

### Inferring signatures of local adaptation

To determine if individual sets of focal loci (identified from GEMMA, bayenv2, and OutFLANK analyses) collectively exhibited elevated signatures of selection acting across multiple loci, we investigated the level of allele frequency covariance among all SNP pairs within each focal set. For instance, to calculate the covariance of allele frequencies across populations between two SNPs, SNP_*i*_ and SNP_*j*_, within a focal set of SNPs associated with a particular phenotype in GEMMA, we used the global minor allele of each SNP, *q*, according to the interpopulation component of linkage disequilibrium,

(1)

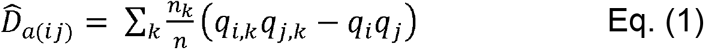

where *n*_*k*_ is the number of individuals in population *k*, *n* is the global population size, *q_i,k_* is the allele frequency of the *i^th^* SNP in population *k, q_i,k_* is the allele frequency of the *j^th^* SNP in population *k*, while *q_i_* and *q_j_* are the respective global allele frequencies of the *i^th^* and *j^th^* SNP across *k* = 6 populations (Storz & Kelly 2008, their Equation 2; Ma *et al*. 2010, their Equation 3). Because we chose the allele to use in comparisons based on global minor allele frequency, all calculations of 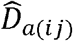 are therefore referenced to the global minor allele haplotype for a pair of SNPs. For populations that experience high levels of gene flow and divergent phenotypic optima due to selection, 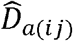 is expected to be positive between allele frequencies of loci conferring a positive effect on the phenotype, negative between those conferring opposite effect, and zero between (conditionally) neutrally loci (eq. [6] in Latta 1998). Because we were not able to discern the direction of effect for alleles within each population (as in e.g., Gompert *et al*. 2015), and to facilitate comparison among analyses, we identified selective signatures by calculating the absolute value of 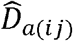 for each locus pair. We also calculated 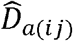 for focal SNPs associated with environmental variables from bayenv2 and those identified as outliers from OutFLANK. In these two cases, we used allele frequencies across all eight populations.

To be able to discern if the level of covariance of allele frequencies among SNPs identified by GEMMA (or another method; hereafter focal SNPs) was greater than that from SNPs randomly chosen from our dataset (i.e., than expected given the data), we first separated all SNPs in the dataset by their expected heterozygosity into bins of 0.01 ranging from 0 to 0.50 (e.g., a SNP with *H*_E_ of (0.000-0.010] would be binned into the first bin, while an *H*_E_ of (0.490-0.500] would be binned into the 50^th^). We then created a set of SNPs from which to take randomized draws by subtracting the focal SNPs from the full set of SNPs. Next, based on the occupancy of heterozygosity bins for a given focal set, we randomly selected remaining SNPs to create a null set. We chose SNPs randomly in this way, 1000 times, each time calculating the absolute value of 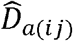 among SNP pairs within each set. From each of these 1000 distributions, we calculated 1000 median absolute 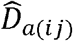 values to create a null distribution for use in comparison to the median absolute 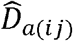 from the focal set of SNPs. If the median 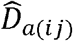 is greater among our focal SNPs than the 95^th^ percentile of the null distribution of 1000 medians, we will conclude that the signature of selection among loci within our focal sets is greater than could have arisen by chance, given the data.

To infer signatures of allele frequency shifts associated to environment, we implemented an approach similar to Equation 1 but instead of estimating 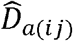 across all populations we estimated 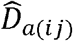 across populations in a pairwise fashion (hereafter *pw*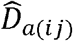) using focal SNPs from a given method. In this case, we calculated global allele frequency (*q_i_* or *q_j_*) based on the frequency of allele *q* across the *k* = 2 populations (*pop*_*l*_ and *pop_m_*) under consideration (where *n_l_* + *n_m_* = *n*). From these estimates, we created a symmetric matrix of *pw*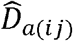 with columns and rows for populations, and distances within the diagonal set to zero. We then implemented Mantel tests (Mantel 1967) using *pw*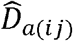 matrices against other population pairwise distance matrices such as geographic distance inferred using great circle distances (km) following Vincenty's method, and Euclidian distance matrices for each of the five phenotypes and 18 environmental variables. Because we chose to take absolute values of *pw*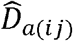 for each locus pair (as with 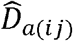) we note that the sign of the correlation coefficient, *r*, from Mantel tests may reflect the opposite directionality for any given SNP pair. Mantel tests were run with 9999 iterations using the skbio package (v0.4.2) in Python. Each environmental or phenotypic value was centered and standardized across populations before calculating Euclidian distances, but not for *pw*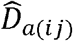 or geographic distance matrices. For each set of focal SNPs associated with phenotype or environment, we also quantified the mean allele frequency differences across populations and compared this to 1000 sets of random SNPs chosen by *H*_E_.

## Results

### SNP filtering and characterization

After calling genotypes, SNPs, and filtering (see results section of Supporting Information), we retained 116,231 imputed SNPs for use as the empirical set in downstream analyses (Table S1). Of these contigs, 107,354 (92.4%) mapped to the *P. lambertiana* reference genome, thus lending authenticity to our sequence data. However, we avoid further discrimination of loci for (proximity to) genic regions until a future genome update with increased curation and density of annotation.

Overall, populations show little genetic structure with plots accounting for less than 1% of the variance in allele frequencies (*F*_plot,total_ = 0.00687; 95% credible interval: 0.0067-0.0070). Of this variation, 56.6% was accounted for by populations (*F*_plot,total_ = 0.00389; 95% CI: 0.0038-0.0040) with the remainder due to plots within populations (*F*_plot,pop_ = 0.00299; 95% CI: 0.0029-0.0031). We found similar patterns among the locus-specific estimates of *F*_ST_ (Figure S1). Moreover, we found no discernable clustering of populations using PCA, respectively accounting for 5.6% and 1.2% of the variance in allele frequencies (Figure S2). To further address applicability of the island model used for calculation of and 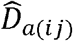, *pw*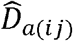 we analyzed population pairwise *F*_ST_ according to Weir & Cockerham (1984) using the hierfstat package in R. Results show little differentiation among populations (mean = 0.005, max = 0.016) with no evidence of isolation by distance (Mantel's *r* = 0.0990, ρ = 0.2310).

### Genotype-environment analyses

To explore the degree of association among environmental variables between populations, we used Mantel tests between Euclidian environmental distance matrices. In most cases, we found significant correlations with many of the edaphic variables measured for this study, as well as between latitude and elevation (*r* = 0.3988, *p* = 0.0490), longitude and annual precipitation (*r* = 0.7145, *p* = 0.0030), and between percent maximum solar radiation and latitude distances (*r* = 0.4629, *p* = 0.0370; Table S2). Additionally, geographic distance among populations was only associated with latitude (*r* = 0.9631, *p* = 0.001), percent maximum solar radiation input (*r* = 0.3992, *p* = 0.0468), and elevation (*r* = 0.4062, *p* = 0.0452), the three of which were correlated environmentally (Table S2), but not to any of the remaining environmental variables (Mantel tests ρ > 0.3131, data not shown).

Through the intersection of the top 0.5% of 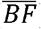 and top 1% of 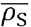, bayenv2 analysis revealed between 14 (CEC) and 157 (GDD-Aug) focal SNPs associated with environment (Table S3). However, when calculating the 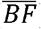 for each SNP, it was never the case that more than two of the five chains produced BF > 1, of which chains with large values were driven primarily by seed number (we used additional seed numbers for a small subset of the data during exploration, data not shown). The range of 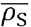 across all focal SNPs across all environments varied from a minimum of 0.138 to a maximum 0.345 (Table S3). Additionally, the focal SNPs identified by bayenv2 displayed a bias towards SNPs with low values of *H*_E_ (Figures S3-S4, see results section of Supporting Information) when compared to the distribution from the full set of SNPs (Figure S5). As such, our environmental associations should be interpreted with caution, as we did not have any SNPs with 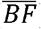 >1 nor do our harmonic mean nonparametric correlations exceed 0.35. Even so, when we compared absolute estimates of 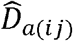 among focal SNPs against the corresponding 1000 null sets of loci, we found that for all focal sets the median 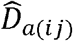 was always greater than the 100^th^ percentile of the null distribution (Figure 2, Table S3). The magnitude of this difference varied across environmental variables, being the smallest for percent clay (1.17x) and largest for annual precipitation (5.10x, Table S3). The percentile of the focal 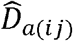 distribution corresponding to the 100^th^ percentile of the associated null set varied across environmental variables as well, reaching just the 3^rd^ percentile for minimum January temperature and the 43^rd^ percentile for percent clay (Table S3). Upon comparison of focal and null sets in both the distribution of single-locus and multilocus *F*_ST_, focal sets were representative of single-locus estimates of the null sets (Figures S6-S7), and generally greater than the distribution of multilocus *F*_ST_ than the null (Figures S8-S9). Single-locus results suggest that many focal SNPs are unlikely outliers for *F*_ST_, while multilocus comparisons exemplify the elevated frequency covariance of SNPs in focal sets. This data demonstrates that for most environmental variables the focal SNPs show higher degrees of 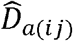 than could be expected, given the data, despite having low 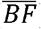 and 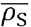.

**Figure 2.**
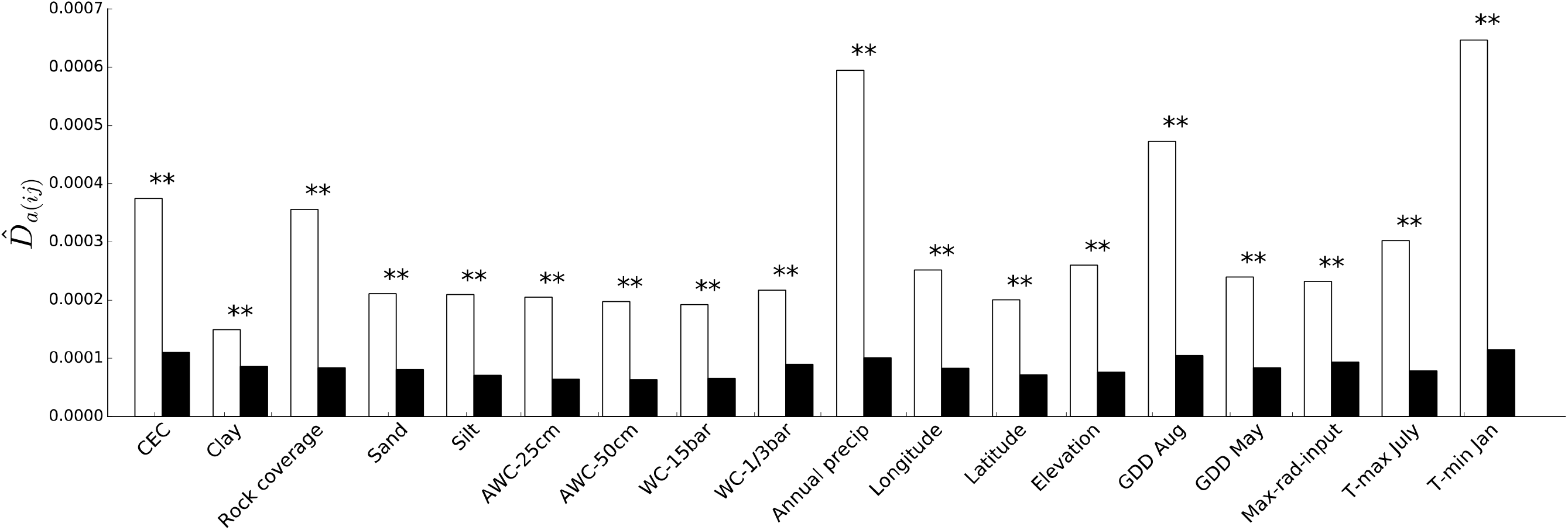
Significantly higher allele frequency covariance (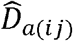) among loci associated to environment by bayenv2 than could be expected to arise, given the data. In white are the median values from 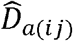 calculated among focal SNPs associated to environment. Black bars display the 95^th^ percentile of the null distribution of median 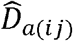. Environmental variables are grouped by those related to soil (CEC through WC-⅓bar) and those related to either climate or geography (Annual precipitation through T-min Jan), with variables related to water availability grouped together in the center of the figure (AWC-25cm through Annual precipitation). All sets of focal loci had median 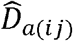 greater than the 100^th^ percentile of the null distribution, as indicated by two stars (**). Environmental variables as in Table 1.

Through the examination of patterns of allele frequency shifts (*pw*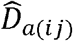) across loci associated with environment we found no significant associations with geographic distance using Mantel tests (*p* > 0.1116). While this suggests the absence of linear allelic clines, it does not necessarily preclude the presence of environmental gradients or correlated patches as suggested by environmental distance associations (Table S2). When we investigated the association between *pw*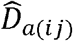 matrices against the eponymous environmental distance matrix, we found significant association for annual precipitation (*r* = 0.7134, *p* = 0.0027), GDD-May (*r* = 0.8480, *p* = 0.0013), longitude (*r* = 0.6522, *p* = 0.0024), percent rock coverage (*r* = 0.5124, *p* = 0.0145), percent sand (*r* = 0.5574, *p* = 0.0046), minimum January temperature (*r* = 0.5791, *p* = 0.0137), and WC-⅓bar (field capacity, r = 0.4806, *p* = 0.0361; Table 2). Additionally, we examined relationships between a particular *pw*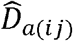 matrix and the 17 remaining environmental distance matrices and found significant associations in an additional 13 comparisons (Table 2), with five of these comparisons having *pw*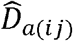 associated with either annual precipitation or longitudinal Euclidian distance. We also observed shifts of alleles associated with longitude or soil water capacity across six of the remaining eight significant associations (Table 2), with the remaining two significant associations among edaphic conditions of sand, silt, or clay. The magnitude of the mean allele frequency difference across populations of focal SNPs was subtle as expected (range 0.018-0.029) and were generally slightly larger than that predicted from random SNPs of the same heterozygosity (Figures S10-S11). Overall, our results indicate that the vast majority of subtle allele frequency shifts among loci associated with environment (*pw*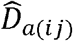) have significant associations related to interpopulation distances of water availability (Table 2).

**Table 2.**
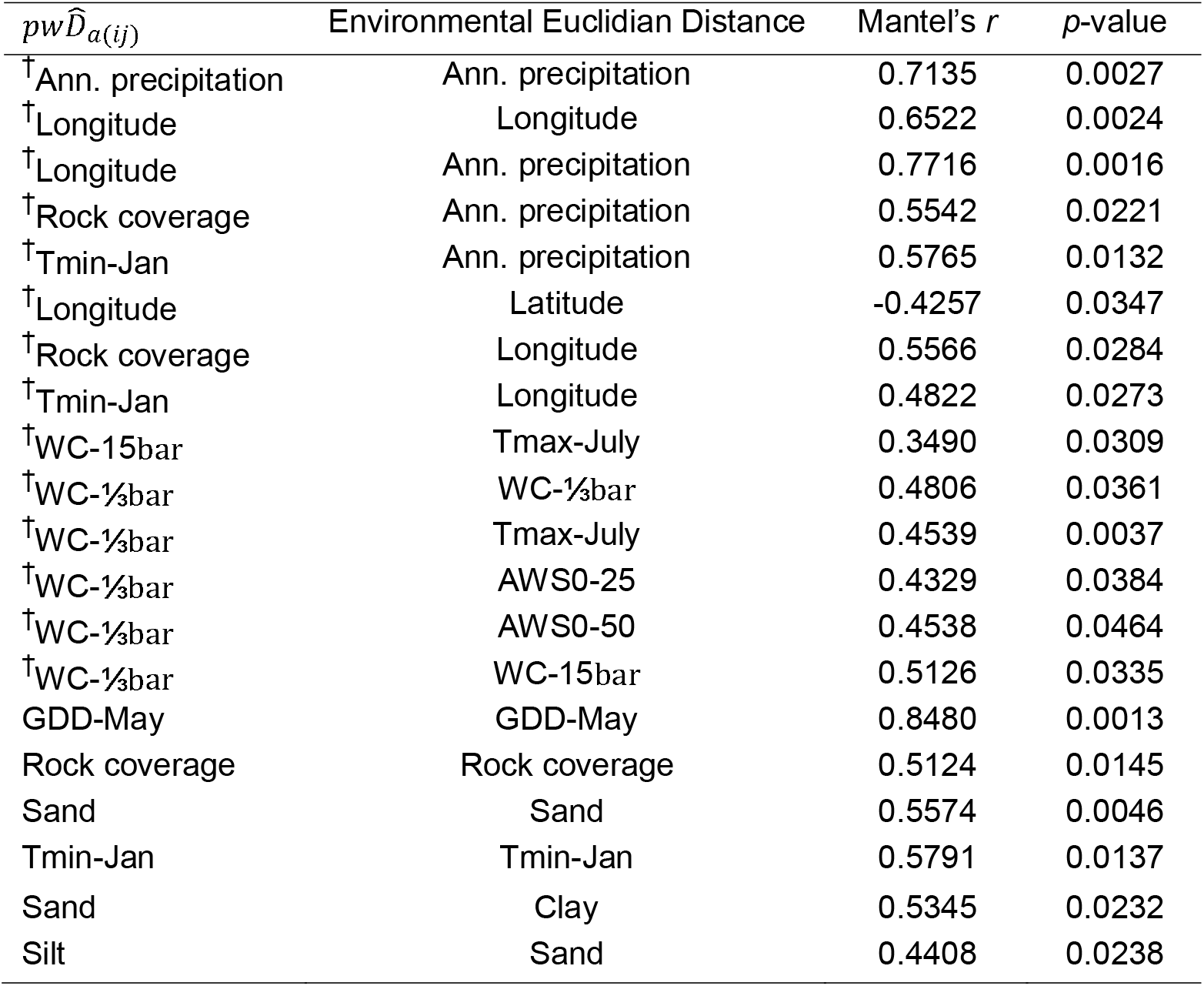
Signatures of allele frequency shifts associated with environmental distance. *Significant Mantel tests (9999 permutations) from comparisons among *pw*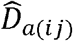 matrices from SNPs associated with environment (first column) against environmental Euclidian distance (second column). Environmental variables as in Table 1.* † indicates a comparison in which at least one variable is water-related, or was associated with annual precipitation.

### Genotype-phenotype analyses

Phenotypic traits were heritable and structured across populations (bud flush *h^2^* = 0.3089 *Q*_ST_ = 0.0156; δ^13^C *h*^2^ = 0.7787, *Q*_ST_ = 0.0427; height *h*^2^ = 0.0608, *Q*_ST_ = 0.0418; N(μg) *h*^2^ = 0.3525, *Q*_ST_ = 0.0191; root:shoot *h*^2^ = 0.3240, *Q*_ST_ = 0.0110; Table S4) and were correlated with environmental variables (both climate and soil) in ways unexplainable by neutral evolutionary forces (Maloney *et al*. in review). Additionally, bud flush and δ^13^C had significant *Q*_ST_ > *F*_ST_ (Maloney *et al*. in review, using the genetic data presented herein). We used a subset of the empirical set of SNPs for use in genotype-phenotype analysis, after filtering we retained 115,632 SNPs. PCA revealed a similar pattern to the empirical set of SNPs (data not shown). Using three significant axes of population structure identified through Tracy-Widom tests, we associated SNPs to phenotypes with BSLMM (Zhou *et al*. 2013) using the top 99.9^th^ and 99.8^th^ percentiles of 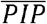 (Figure 4, Table S5, Figure S12). From observations of density and trace plots, we concluded that the posterior distributions across chains were converging (not shown). The *H*_E_ of focal loci were generally representative of the empirical set (Figures S13-S14).

Overall, the genetic variance of SNPs included in the polygenic model explained between 14.4% [N(μg)] and 37.6% (root:shoot) of the variance in the phenotypes measured in our study (*PVE*, Figure 3, Table S4). For many of the measured phenotypes, a considerable proportion of the narrow sense heritability estimated previously was therefore accounted for in the estimates of *PVE* (Table S4), as should be the case with sufficient genetic sampling (Gompert *et al*. 2016). Interestingly, in the case of height, *PVE* exceeded the upper confidence interval of the estimated *h*^2^ (Table S4).

**Figure 3.**
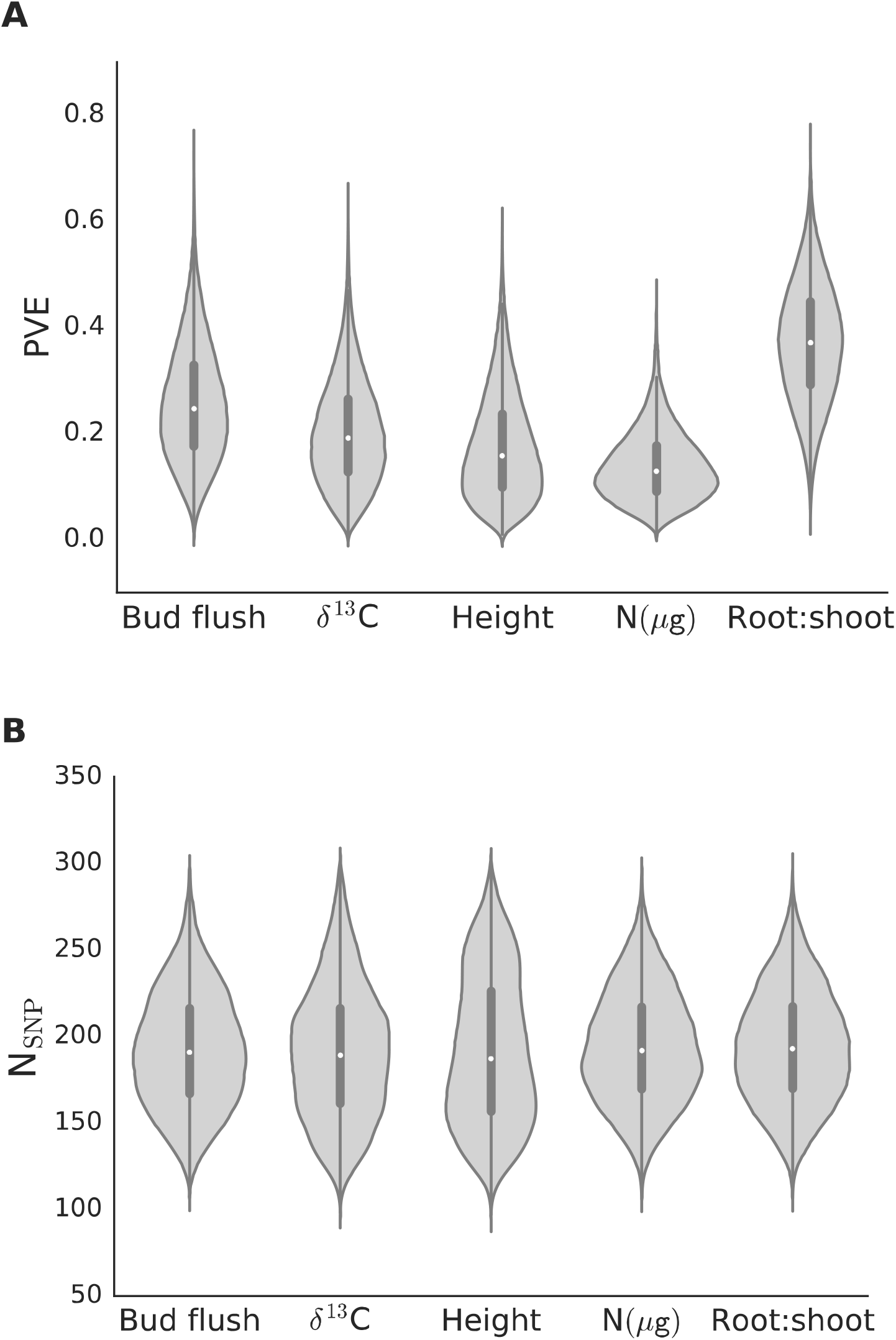
Violin plots for the kernel density estimator of the posterior distributions (light grey) taken from Bayesian sparse linear mixed models (BSLMM) executed in GEMMA for (A) the proportion of variance explained by SNPs included in the model (PVE) and (B) the number of SNPs underlying the phenotypic trait (N_SNP_). Priors for N_SNP_ and PVE were [1,300] and [0.01,0.9], respectively. Dark grey vertical bars display the first through third interquartile range with the median represented by the white dot.

To acquire estimates of *PVE* from the identification of loci with large effects on phenotype, we conducted single-locus association using univariate linear mixed models implemented in GEMMA (see Supporting Information, Table S6). Across all phenotypes, there were no loci that exceeded the adjusted threshold for inclusion calculated from *q*-values with an FDR of 0.05 (Storey *et al*. 2015; v2.4.2), with the minimum q-value across SNPs within phenotypes ranging between 0.2046 (δ^13^C) and 0.9999 [N(μg)] (Table S6). Except for root:shoot biomass, the maximum likelihood estimates of *PVE* differed drastically from the estimates from BSLMM, with *PVE* never exceeding 1.08e-06 suggesting that a larger proportion of the heritable genetic variation for the traits measured here is explained by multiple SNPs than by individual SNPs alone. Finally, to determine if LMM loci near the threshold were captured by the BSLMM for a particular phenotype, we isolated the loci from univariate LMM above a reduced threshold of -*ln(pw_ald_*) ≥ 10 (see Figure S15, Supporting Information). By this reduced threshold we identified one unique locus for both bud flush and N(μg), four unique loci for both height and root:shoot biomass, and five unique loci for δ^13^C (15 unique loci overall). We examined the focal loci sets identified from the 99.9^th^ percentile of 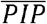 in BSLMM for these LMM reduced-threshold loci and found 1 of the 4 LMM loci for both root:shoot biomass and height, and 2 of the 5 loci for δ^13^C When we assessed the set of loci in the 99.8^th^ percentile of BSLMM 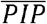, we recovered all LMM reduced-threshold loci for bud flush and N(μg) (n = 1), 1 of 4 loci for root:shoot biomass, 3 of 4 loci for height, and 3 of 5 loci for δ^13^C.

To determine if focal loci associated with phenotype by BSLMM exhibited evidence of selection, we estimated allele frequency covariance (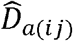) among focal SNPs and compared these estimates to 1000 null sets of SNPs. We found evidence for elevated covariance among the 99.9^th^ percentile of 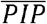 loci associated with bud flush and root:shoot biomass (Figure 4a, Table S5), with the latter exceeding the 100^th^ percentile of the random distribution. To consider larger numbers of loci representative of the number of underlying loci estimated by BSLMM, we also isolated SNPs from the top 99.8^th^ percentile of 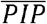. In these sets, we found evidence for elevated signatures of selection acting across multiple loci for all phenotypes except for height, which did not produce a focal median 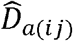 greater than the 95^th^ percentile of null distribution of 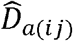 (Figure 4b, Table S5). When focal and null sets of SNPs were compared, focal sets were representative of single-locus (Figure S16) and multilocus (Figure S17) *F*_ST_ estimates of the null sets.

**Figure 4.**
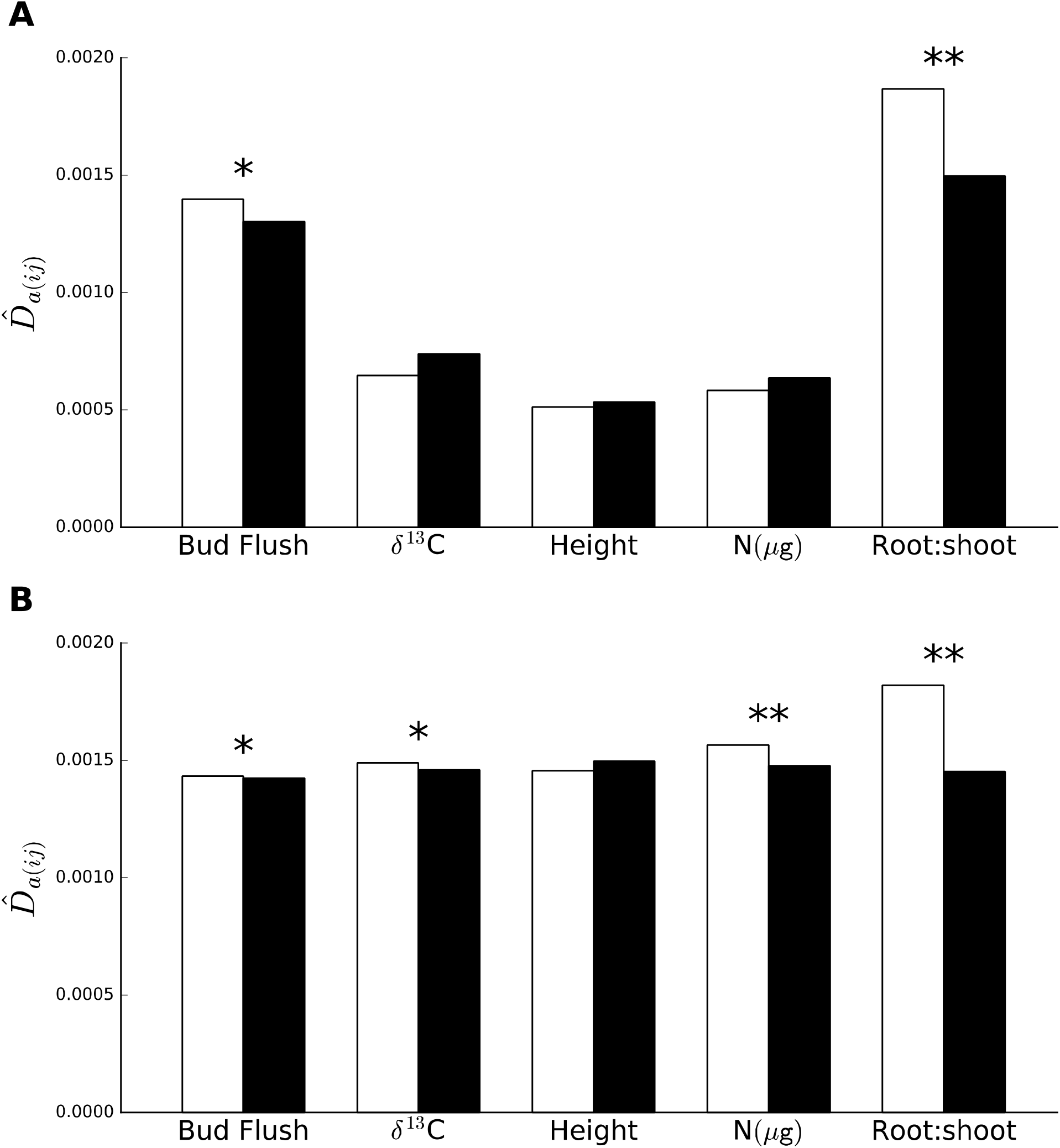
Allele frequency covariance (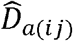) among loci associated to phenotype by GEMMA. In white are the median values from 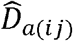 calculated among focal SNPs associated to phenotype. Black bars display the 95^th^ percentile of the null distribution of median 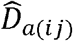 (A) SNPs identified in the top 99.9^th^ percentile of 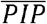, (B) SNPs identified in the top 99.8^th^ percentile of 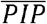. One star (*) indicates that the median focal 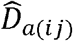 was greater than the 95^th^ percentile of the null distribution, whereas two stars (**) indicate that the focal median 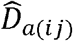 was greater than the 100^h^ percentile of the null distribution.

To identify signatures of allele frequency shifts among focal loci associated with phenotype (*pw*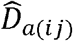), we ran Mantel tests of *pw*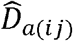 matrices against geographic distance and environmental Euclidian distance matrices. When considering SNPs identified by the 99.9^th^ percentile of 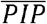, we see substantial evidence for allele frequency shifts of loci associated with bud flush to Euclidian distances of GDD-May, GDD-Aug, percent maximum radiation input, and minimum January temperature (Table 3). Additionally, when we consider the 99.8^th^ percentile of 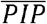, we show evidence for allele frequency shifts among loci associated with bud flush, height, and δ^13^C to Euclidian distances of annual precipitation, as well as for N(μg) loci to elevation, and bud flush loci to both longitude (a correlate of annual precipitation) and to percent maximum radiation input (as in the 99.9^th^ 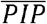 set). The strong signal from bud flush and water-related variable in Table 3 is intriguing, as bud flush and δ^13^C were the only two phenotypic traits to have significantly larger *Q*_ST_ than *F*_ST_ (Maloney *et al*. in review). The magnitude of the mean focal allele frequency differences across populations were subtle as expected (range: 0.054-0.087) and representative of unassociated SNPs of similar *H*_E_ (Figure S18).

**Table 3.** Signatures of allele frequency shifts associated with environmental distance. *Significant Mantel tests (9999 permutations) from comparisons among allele frequency shifts (*pw*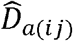) of SNPs associated with phenotype against environmental Euclidian distance. Selection Criterion refers to the process used to identify SNPs associated with phenotype.* † indicates a comparison in which at least one variable is water-related, or was associated to annual precipitation.

| Selection criterion | $pw\hat{D}_{a(ij)}$ | Environmental Euclidian Distance | Mantel's $r$ | $p$-value |
| --- | --- | --- | --- | --- |
| 99.9<sup>th</sup> $\overline{PIP}$ | Bud flush | GDD August | 0.5804 | 0.0181 |
|  | Bud flush | GDD May | -0.5190 | 0.0458 |
|  | Bud flush | Max. radiation input | -0.5486 | 0.0482 |
| | Bud flush | $T_{\min}$ January | 0.6984 | 0.0191 |
| 99.8<sup>th</sup> $\overline{PIP}$ | <sup>†</sup>Bud flush | Annual precipitation | 0.4309 | 0.0140 |
|  | <sup>†</sup>Bud flush | Longitude | 0.5532 | 0.0405 |
|  | Bud flush | Max. radiation input | -0.6312 | 0.0127 |
| | N(μg) | Elevation | 0.5334 | 0.0246 |
|  | <sup>†</sup>Height | Annual precipitation | 0.7210 | 0.0320 |
| | <sup>†</sup>$\delta^{13}\text{C}$ | Annual precipitation | 0.5952 | 0.0195 |

### *F*_ST_ outlier analysis

From the empirical set of SNPs, OutFLANK analysis revealed 110 focal loci as outliers for *F*′_ST_ (range: 0.069-0.118). Expected heterozygosity values among the outlier SNPs (Figure S19) varied across the distribution from the full set of SNPs (Figure S5). Upon analysis of patterns of covariance (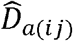) among the OutFLANK focal SNPs, we found that the median focal 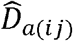 (6.08e-03) was 10.6x greater than the 100^th^ percentile of the null distribution of 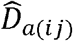 (5.74e-04). Moreover, the maximum median 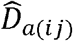 from null sets corresponded to just the 12^th^ percentile of the focal 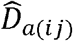 values, suggesting that the majority of SNPs within the focal set showed higher levels of covariance among other outliers than expected by chance. However, when we analyzed these outlier SNPs for signatures of allele frequency shifts (*pw*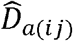) we found no significant associations with geographic or environmental distances.

### Intersection of SNPs within and across methods

We examined overlap of focal SNPs among the various methods employed in this study (Table S7). While there was considerable overlap of loci found between methods, overall there was more overlap of loci associated with multiple phenotypes or with multiple environments than found across methods. For sets of loci associated with environmental variables, there was a considerable number of loci that were found to overlap among comparisons. This seemed to be driven by the correlations among soil properties, for when ordered by the number of loci within the intersection, 12 of the top 15 comparisons were among edaphic conditions. Additionally, climate-related variables relating to maximum radiation input, degree growing days, minimum and maximum temperature generally shared loci among soil variables relating to water availability while annual precipitation shared 18 loci with longitude, among other variables. Very few of the loci identified by bayenv2 would have been detected through conventional *F*_ST_ outlier approaches (Figures S20-S21). Even so, OutFLANK captured between 1-3 (n = 18) of the loci identified across 10/18 environmental associations from bayenv2 (many of which were water-related, including annual precipitation), and captured 4 of the loci identified in the 99.8^th^ percentile of 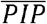, but not for any of the loci identified from the reduced threshold of LMM. Among loci associated with phenotype (99.8^th^ 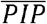), there were between one and three loci which were found in the intersection among pairwise phenotypic comparisons, yet none of these overlap loci were those identified from LMM. Finally, 15 loci associated with environment overlapped with the 99.8^th^ percentile of 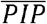 (including two between δ^13^C and longitude, a correlate of annual precipitation) while environmental associations did not capture any of the reduced-threshold loci from univariate LMM (Table S7).

## Discussion

The spatial extent of local adaptation, particularly in conifers, has generally been investigated at regional scales (Neale & Savolainen 2004; Savolainen *et al*. 2007; Ćalić *et al*. 2016). While informative for range-wide inference, management and conservation agencies are often limited to local scales spanning only tens to several hundreds of square kilometers. While there is an expectation that high gene flow (i.e., migration load) exhibited by many conifers can lead to swamping of adaptive alleles, there is mounting empirical evidence that adaptation to the environment can still occur at relatively fine spatial scales (Mitton 1989; 1999; Budde *et al*. 2014; Csilléry *et al*. 2014; Vizcaíno *et al*. 2014; Eckert *et al*. 2015). Thus, studies which investigate adaptation at scales amenable to management may be of relatively greater importance (especially for endangered and threatened species) to reforestation applications such as those carried out through seed sourcing (*sensu* McLane & Aitken 2012) and replanting efforts. Previously, we provided evidence that measured fitness-related phenotypes are heritable, that population explains a significant proportion of phenotypic variation, and *Q*_ST_ was significantly greater than *F*_ST_ for bud flush and ô^13^C (Maloney *et al*. in review). Here, our genetic analyses indicate selective pressures of *P. albicaulis* are likely driven by water availability (e.g., precipitation gradients) as well as interactions and correlates of soil properties, which lends replicative support to both theoretical and empirical predictions for the patterns of loci underlying quantitative traits undergoing selection with gene flow. For instance, Ma *et al*. (2010) assessed evidence for diversifying selection within European aspen (*Populus tremula* L.) across 23 candidate genes of the photoperiodic pathway using the covariance of allelic effects among loci, albeit across a geographic region of Sweden spanning 10 latitudinal degrees. From this candidate set, they identified high degrees of covariance among phenotypic effects as predicted from theory (Latta 1998), despite minimal allele frequency differentiation among sampled populations. More recently, Csilléry *et al*. (2014) assessed 53 climate-related candidate genes within European beech (*Fagus sylvatica* L.) providing evidence that covariance among loci is attributable to epistatic selection (*sensu* Ohta 1982) across fine spatial scales of less than 100km^2^. While varying across spatial scales, these studies replicate evidence for a signal of local adaptation in trees through elevated among-population linkage disequilibrium between adaptive loci.

### Standing genetic variation for fitness-related traits

The populations under study appear to have extensive gene flow, recent divergence, or both. Variation of allele frequencies among populations accounts for less than 1 of the variance observed, which was less than that found for *P. lambertiana* populations within the LTB (Eckert *et al*. 2015), or among isozymes sampled from populations across the Northern *P. albicaulis* range (Krakowski *et al*. 2003). Inspection of PCs showed no distinctive clustering of populations (Figure S2) while population pairwise *F*_ST_ did not exceed 0.016 and a test for isolation by distance was not significant. Consequently, the mean allele frequency differences between populations for focal SNPs were also subtle. Importantly, the majority of the populations under study exhibited significant, heritable genetic variation underlying the measured phenotypic traits. Biologically, such a pattern of extensive sharing of alleles across populations has likely resulted from a combination of long-distance pollen movement, and seed dispersal by Clark's nutcracker (*Nucifraga columbiana* Wilson) which is known to disperse seeds at distances similar to those between our sampled populations (Tomback 1982; Richardson *et al*. 2002 and references therein). Given this pattern of structure, the island model with symmetric migration used to describe the interpopulation component of linkage disequilibrium among loci (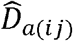) and allele frequency shifts (*pw*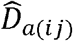) is likely suitable to investigate our dataset for signatures of selection across multiple loci.

While bayenv2 did not identify any loci strongly associated with environment, as given from small values of Bayes factors (all 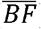 < 1.0, Table S3), there is a strong biological signal for adaptation to soil water availability in our dataset (discussed below), evidence that other white pines within the LTB are also being structured by precipitation differences among populations (Eckert *et al*. 2015), and elevated signals of selection among focal loci which would be unlikely to arise, given the data. Thus, it seems unlikely that the focal sets of SNPs are driven solely by false positives. However, if the majority of loci across bayenv2 are not an artifact of the method (i.e., are within or in linkage with causative sites), one possible explanation for elevated covariance among focal loci is that the structure of environmental variables across populations captured variation for unmeasured phenotypic traits which were largely representative of total lifetime fitness (Schoville *et al*. 2012). Structure of unmeasured fitness-related traits may also explain the high covariance of OutFLANK loci. Future work could provide validation through functional analyses of loci or from similar patterns found in other systems.

### Water availability as a driver of local adaptation

The strongest signal for local adaptation among *P. albicaulis* populations of the LTB came from evidence of adaptation to soil water availability (Figures 2 and 4; Tables 2–3). Indeed, water availability is a critical component shaping standing variation across plant taxa (Vicente-Serrano *et al*. 2013), including the distributions of tree species in general (van Mantgem *et al*. 2009; Allen 2010), and southern populations of *P. albicaulis* specifically (Bower & Aitken 2008; Chang *et al*. 2014). During the Pleistocene-Holocene transition (10,000-12,000yr BP), shifts from mesic to xeric conditions caused proximal *P. albicaulis* populations of the Western Great Basin (~50km distant) to shift from 1380m in elevation to their current position about 1100m down-slope (Nowak *et al*. 1994; *cf.* Table 1). Such shifts in climate and local edaphic conditions in the last 10,000yr may in part explain recent (relative to 4*N*_e_ generations), and ongoing, selective pressures on *P. albicaulis* populations of the LTB. Because of climatic constraints imposed on the southern range of *P. albicaulis*, phenotypic traits affected by precipitation, soil water availability, or soil water capacity likely have fitness-related consequences for this species. Additionally, with climatic models predicting warmer temperatures, reduced snow accumulation, and earlier spring melt across the western USA, it is likely that *P. albicaulis* populations of the Sierra Nevada will continue to face selective pressures of this kind. Even so, many of *P. albicaulis* populations of the LTB exhibit substantial genetic variation for the fitness-related traits measured, suggesting that the majority of ongoing adaptation within the LTB will likely be unconstrained by the lack of genetic variation. Instead, other biotic factors (e.g., white pine-blister rust infection) that can lead to negative population growth rates may be of more immediate concern (see Maloney *et al*. 2012).

### Implications to whitebark management

As pointed out by McLane & Aitken (2012), distribution models of many species predict climate niches to shift in the upcoming century, leaving great uncertainty that tree species in particular will be able to track suitable environments through natural migration and establishment, as the rate of many of these geographic shifts would far exceed observed post-glacial rates of migration (Davis & Shaw 2001; McLachlan *et al*. 2005). Exacerbating this issue, the presence of *C. ribicola*, and climate-driven outbreaks of mountain pine beetle *(D. ponderosae)* create further challenges to the conservation of *P. albicaulis* (Tomback & Achuff 2010; Mahalovich & Stritch 2013). Without intervention, such cases could lead to population collapse, extirpation, or extinction. As such, assisted gene flow, replanting, or restoration efforts will need to continue to take current and future selective pressures into account (e.g., genetic variation, resistance to *C. ribicola*, etc.), as has generally been the standard of practice (Keane *et al*. 2012; Maloney *et al*. 2012; McLane & Aitken 2012).

At the same time, the choice of seed source will also need to take into account local adaptation at fine spatial scales. While a small proportion of the neutral genetic variation (*F*_ST_) is found among most tree populations (often less than 5%, Neale & Savolainen 2004), this does not necessarily mean that seeds from within an established seed zone will be optimal for any given constituent environment, particularly if the seed zone exhibits high environmental heterogeneity (such as in montane regions), or if the seed zone is relatively broad compared to these environmental gradients, as is the case in California (see Buck *et al*. 1970). Weak neutral genetic differentiation can be misleading in this way, as polygenic traits influenced by selective processes in the face of gene flow may lead to divergent local adaptation through the covariance of alleles among populations without the buildup of substantial genetic differentiation at any given locus. Particularly in cases where there is evidence of local adaptation, and when ethical (McLachlan *et al*. 2007), appropriate (considering e.g., ecological or demographic factors), or plausible, seed source should come from local sources (i.e., relative to scales of geography and environmental gradients) to maximize adaptive potential (McKay & Latta 2002). This is particularly important when maximal fitness (e.g., reproductive output) is a priority for established trees, as while many genotypes may survive, realized phenotypes related to fitness may be suboptimal for a given environment. In cases where local seed source is not plausible, perhaps due to (inbreeding through) isolation, or with prohibitively small population sizes, sources likely to perform well are also likely to come from highly correlated environments, particularly if the populations are not highly diverged. With this taken into consideration, management may be able to prioritize populations for restoration through estimates of trait heritability (the prerequisite for adaptation), as estimates of genetic variation alone may be misleading. For instance, heritability for δ^13^C was near zero for the Rifle Peak population, as was the case for root:shoot ratio in Freel Peak and Little Roundtop, δ^13^N in Rifle Peak, and height for four of six populations. While this could mean the presence of recent, strong natural selection, or, conversely, that these traits do not convey an overwhelming adaptive advantage in these populations, candidate traits with substantial evidence for contributing to total lifetime fitness should be monitored nonetheless. In appropriate cases, introducing compatible variation would stand to improve adaptive potential in such populations. While there is no one specific solution to conserve whitebark populations across its range, taking into consideration fine-scale local adaptation in addition to established strategies will likely aid in such endeavors.

### Concluding remarks, limitations, and future work

The results reported here suggest that focal loci collectively show elevated allele frequency covariance (a signal expected between loci undergoing selection with gene flow) across multiple loci than could have been expected to arise from the data by chance, and that interpopulation levels of allele frequency covariance are often associated with interpopulation distances of soil water availability. Our results further explain a considerable proportion (*PVE*) of the additive genetic variation (*h*^2^) of the quantitative traits under study from a polygenic perspective, as should be the case with sufficient genetic sampling. Thus, we can posit that the general mode of adaptation for *P. albicaulis* across the LTB is facilitated by selection on standing levels of genetic variation that is extensively shared throughout the basin and likely improves performance in early life stages. Finally, if soil and climatic variables continue to influence the extant populations within the LTB as evidenced from our analyses, it is likely that these variables will continue to be important to the long-term success of this threatened keystone species.

While we described associations among genotype, phenotype, and environment that reflect strong evidence for adaptive responses of *P. albicaulis* populations to the environment, we acknowledge several shortcomings. First, our study design was limited in statistical power which could have been improved by increasing the number of individuals sampled, the total number of populations, or both, given an ideal sampling regime (Lotterhos & Whitlock 2015) which together would have further facilitated other methodologies for uncovering evidence of polygenic local adaptation (e.g., Berg & Coop 2014). Second, while we measured fitness-related traits among seedlings of a species whose lifespan differs by several orders of magnitude, establishment success is one of the primary factors influencing dynamics of forest populations, and early life stages of plants have been shown to be a major component of total lifetime fitness (Postma & Âgren 2016). Third, much of the statistical signal for the association of allele frequency shifts to environment would be lost with correction for multiple tests yet we leverage the fact that, of the few significant *pw*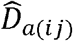 associations, the majority were related to δ^13^C annual precipitation, its correlate of longitude, or measures of soil water availability, an outcome highly unlikely by chance alone. Fourth, while we provide evidence for statistical signals predicted by theory, our methodology limited us from making conclusions regarding local adaptation *sensu stricto* as we utilized just a single common garden without reciprocal transplants and were unable to quantify functional differences of putative loci among populations. Reciprocal transplants would have allowed us to differentiate pleiotropic effects and facilitate direct measures of fitness through survival and growth across environments. Finally, a more fully curated, well-annotated genome assembly and accompanying linkage map would have aided in the detection of physical linkage among SNPs, proximity to genomic regions of estimated effect, and detection of false positives. For instance, the *P. lambertiana* genome used to judge authenticity of sequence data does not yet have the density of annotation needed to draw inferences on the causative sites likely within or linked to the loci described here, as its assembly and curation are still ongoing. Because of this, we cannot confidently estimate the proportion of the polymorphism due to coding and non-coding sites nor conclude that the linkage inferred for focal loci are not an artifact due to distant linkage with causative sites of larger effect. For this artifact to be true, however, our results would indicate that many of the loci in focal sets of SNPs would all have to be linked to the same smaller number of larger-effect loci, or that many large-effect loci underlie the trait, both unlikely outcomes given the expectations of quantitative traits and the coverage from ddRADseq methods found in other white pines (e.g., Friedline *et al*. 2015). Even so, while we may not have identified all causative sites, linked sites can still maintain signals of evolutionary processes (McVean 2007). Moreover, signatures of selection presented here are unlikely to have arisen from our dataset by chance alone (as evidenced from null models) and our inferences were synthesized from prevailing signals across multiple phenotypic, environmental, and taxonomic lines of evidence, as opposed to inferences based on single variables or tests. Future work could address these shortcomings and lead to the corroboration of our results, particularly in describing patterns exhibited by underlying loci in similar systems.

## Acknowledgements

We thank Annette Delfino Mix, Camille Jensen, Tom Burt, and Randi Famula for field, common garden, and lab assistance. Additionally, we thank David Fournier, Joey Keely, Kurt Teuber (USDA Forest Service – LTBMU), Roland Shaw (Nevada Division of Forestry), Bill Champion (Nevada State Parks), Woody Loftis (USDA NRCS) for site information and permission to work on Federal and State lands, the VCU CHiPC for computational resources, Jennifer Ciminelli for GIS tips, and Lindsay Miles who helped improve this manuscript. This work was supported by the Southern Nevada Public Lands Management Act – Rounds 7 and 10, sponsored by the USDA, Forest Service, Pacific Southwest Research Station, Albany, CA. We also thank anonymous reviewers whose suggestions greatly contributed to this manuscript.

## Data Accessibility

Sequence data is deposited in the short read archive of the National Center for Bio-technology Information (project number: TBD). Scripts used in analyses can be found in IPython notebook format (Pérez and Granger 2007) at https://github.com/brandonlind/whitebarkpine.

## Author Contributions

PEM, AJE, DBN, DRV, and JLW conceived the study. PEM oversaw sample collection and common garden maintenance. BML wrote the manuscript and carried out analysis of data, with contributions from CJF and AJE. All authors contributed to the editing of the manuscript.

